# Tractography-based automated identification of the retinogeniculate visual pathway with novel microstructure-informed supervised contrastive learning

**DOI:** 10.1101/2024.01.03.574115

**Authors:** Sipei Li, Wei Zhang, Shun Yao, Jianzhong He, Ce Zhu, Jingjing Gao, Tengfei Xue, Guoqiang Xie, Yuqian Chen, Erickson F. Torio, Yuanjing Feng, Dhiego CA Bastos, Yogesh Rathi, Nikos Makris, Ron Kikinis, Wenya Linda Bi, Alexandra J Golby, Lauren J O’Donnell, Fan Zhang

## Abstract

The retinogeniculate visual pathway (RGVP) is responsible for carrying visual information from the retina to the lateral geniculate nucleus. Identification and visualization of the RGVP are important in studying the anatomy of the visual system and can inform the treatment of related brain diseases. Diffusion MRI (dMRI) tractography is an advanced imaging method that uniquely enables *in vivo* mapping of the 3D trajectory of the RGVP. Currently, identification of the RGVP from tractography data relies on expert (manual) selection of tractography streamlines, which is time-consuming, has high clinical and expert labor costs, and is affected by inter-observer variability. In this paper, we present a novel deep learning framework, *DeepRGVP*, to enable fast and accurate identification of the RGVP from dMRI tractography data. We design a novel microstructure-informed supervised contrastive learning method that leverages both streamline label and tissue microstructure information to determine positive and negative pairs. We propose a simple and successful streamline-level data augmentation method to address highly imbalanced training data, where the number of RGVP streamlines is much lower than that of non-RGVP streamlines. We perform comparisons with several state-of-the-art deep learning methods that were designed for tractography parcellation, and we show superior RGVP identification results using DeepRGVP. In addition, we demonstrate a good generalizability of DeepRGVP to dMRI tractography data from neurosurgical patients with pituitary tumors and we show DeepRGVP can successfully identify RGVPs despite the effect of lesions affecting the RGVPs. Overall, our study shows the high potential of using deep learning to automatically identify the RGVP.

## 1. INTRODUCTION

The retinogeniculate visual pathway (RGVP) is responsible for carrying visual information from the retina to the lateral geniculate nucleus (LGN) (Chacko, 1948). It consists of three anatomical segments, including the optic nerve, the optic chiasm, and the optic tract (J. He et al., 2021). The RGVP is affected in many diseases, including pituitary tumors (Laws et al., 1977), optic neuritis (Beck et al., 2003), optic nerve sheath meningiomas (Schick et al., 2004), optic gliomas (Hales et al., 2018), and many others (Attyé et al., 2018; Pisa et al., 2022). In particular, pituitary tumors notably influence the RGVP, because of their close proximity to the optic nerves and chiasm, and can lead to neurological complications such as vision impairment, headaches, and nausea (Gittleman et al., 2014; Ho et al., 2015; Rucker & Kernohan, 1954). Identification and visualization of the RGVP are important in studying the anatomy of the visual system (Wichmann & Müller-Forell, 2004) and may inform treatment of brain diseases such as lesions intrinsic or extrinsic to the pathway (Ma et al., 2016).

**Figure 1.**
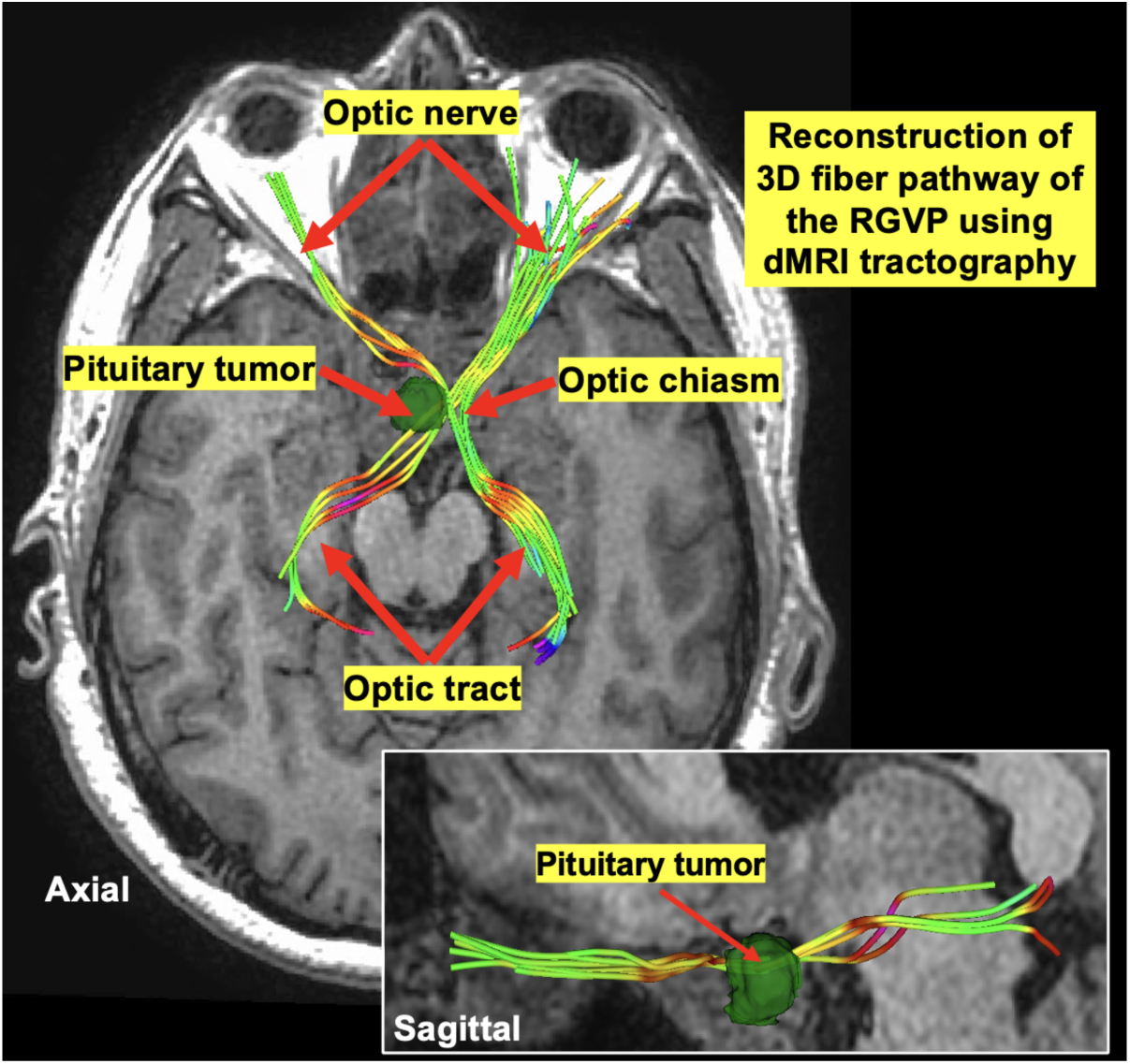
Mapping of the retinogeniculate visual pathway (RGVP) in a retrospective pituitary adenoma patient using dMRI tractography. The 3D fiber pathway of the RGVP is reconstructed consisting of three anatomical segments, i.e., the optic nerves, the optic chiasm, and the optic tracts. The tumor (in green) causes displacement of RGVP fibers near the optic chiasm region.

Diffusion MRI (dMRI) tractography is an advanced imaging method (Basser et al., 2000) that uniquely enables *in vivo* reconstruction of the 3D streamline trajectory of the RGVP in a non-invasive way. Unlike the widely used T1-weighted (T1w) and T2-weighted (T2w) MRI or other advanced imaging techniques, e.g. steady-state free precession (SSFP), that can only indicate whether the RGVP is present at a particular location, dMRI tractography can assess the 3D fiber pathway of the RGVP and the continuity of its subdivisions. In the context of surgical planning, dMRI tractography allows neurosurgeons to visualize the relationship of the RGVP and its subdivisions to the surrounding structures. In this way, it allows neurosurgeons to evaluate the proximity of the planned surgical path. As shown in Figure 1, dMRI tractography can enable the visualization of the 3D pathway of the RGVP and its relation to the tumor. Due to this unique property, dMRI tractography has been used successfully in presurgical cranial nerve (CN) mapping (Roundy et al., 2012) for making optimal surgical plans, improving the possibility of preserving CNs and contributing to improved quality of life (Churi et al., 2019; Song et al., 2016).

Many tractography-based studies have shown successful mapping of the RGVP for clinical and research purposes (Altıntaş et al., 2017; Ather et al., 2019; Hofer et al., 2010; Panesar et al., 2019; Yoshino et al., 2016). Currently, identification of the RGVP from tractography mainly relies on expert selection, where streamlines are selected based on whether they end in and/or pass through regions of interest (ROIs) drawn by experts (Oishi et al., 2008). ROI-based RGVP selection, however, is time-consuming, is inefficient with high clinical and expert labor costs, and is also affected by inter-observer variability depending on the experience of experts. Therefore, there is a high need for computational methods to enable automated identification of the RGVP. Recently, there have been studies for automated identification of the RGVP as well as other cranial nerves (Zeng et al., 2023; Zhang, Xie, et al., 2020) using traditional machine-learning techniques. In recent years, deep-learning-based methods have been demonstrated to be highly successful in tractography parcellation for automated identification of white matter fiber tracts in the cerebrum (Y. Chen et al., 2023; Wasserthal, Jakob, Peter Neher, and Klaus H. Maier-Hein., 2018; H. Xu et al., 2019; Zhang, Karayumak, et al., 2020). To date, there have been no deep-learning-based methods for RGVP identification using tractography in surgical patient data with lesions affecting the RGVP.

Recent advances in deep learning provide a promising approach to enable accurate and fast identification of the RGVP. However, there are two key challenges. First, current deep-learning-based tractography parcellation methods mostly use streamline geometric features extracted based on the streamline point spatial coordinates (Gupta et al., 2017; Lam et al., 2018; Xue et al., 2023a; H. Xu et al., 2019), e.g., RAS (Right, Anterior, Superior) coordinates. While this spatial information is effective in differentiating white matter fiber tracts that have large shape and position dissimilarities, there can be very small geometric differences between RGVP streamlines and nearby non-RGVP streamlines that may belong to other structures (e.g., orbital muscles) or result from false positive streamline tracking due to partial voluming and low SNR issues (Carrozzi et al., 2023; J. He et al., 2021; Hofer et al., 2010). Thus, only using geometric features can be ineffective for RGVP identification. In tractography data, various features can be computed, including not only streamline geometric features but also diffusion microstructure measures like fractional anisotropy (FA) (Zhang, Daducci, et al., 2022). We hypothesize that including such microstructure features can improve identification of the RGVP. Our rationale is that streamlines representing similar anatomical structures should not only have a similar geometric trajectory but also should have a very similar FA value. A second challenge is that there can be a highly imbalanced streamline sample distribution between RGVP and non-RGVP streamlines (our data shows a ratio of 1:8 between RGVP and non-RGVP streamlines; see Sec. 2.1). Imbalanced training data is a well-known challenge in deep learning and limits network generalization for small-size sample categories (Johnson & Khoshgoftaar, 2019). Data augmentation (DA) is one effective solution to resolve this challenge. In related work, several tractography studies have performed DA by generating synthetic data samples, e.g., duplicating and/or adding noise to the existing tractography data (Benou & Raviv, 2019; Gupta et al., 2017). Our recent study proposed a subject-level tractography DA strategy that generates multiple new datasets by downsampling each subject’s tractography data, without creating synthetic data (Zhang, Xue, et al., 2022). Inspired by this work, we propose a new streamline-level DA method that uses random subsampling to increase the sample size of RGVP streamlines. We hypothesize that this method can generate a balanced dataset for improved model training.

In light of the above, this study presents a novel deep learning framework, namely *DeepRGVP*, for automated identification of the RGVP using dMRI tractography. In this study, our contributions are as follows. First, we present what we believe is the first deep-learning approach that enables fast and accurate RGVP identification. The method is based on our Superficial White Matter Analysis (SupWMA) framework (Xue et al., 2023a) that uses a point-cloud-based network (Qi et al., 2017) and supervised contrastive learning (SCL) (Khosla et al., 2020) for the classification of superficial white matter streamlines. Second, we design a microstructure-informed SCL method that leverages both streamline label (RGVP and non-RGVP) and tissue microstructure (FA) information to determine positive and negative pairs to enable contrastive learning. In this way, the learned global feature may have the ability to better classify streamlines with similar trajectories but different classes. Third, we design a simple and successful streamline-level data augmentation method to increase the training sample size. In this way, we can effectively address the imbalanced training data where the number of non-RGVP streamlines is much more than that of the RGVP streamlines. Fourth, we demonstrate successful generalization of DeepRGVP to clinically typical dMRI data acquired from pituitary tumor patients, even though DeepRGVP is trained using high-quality dMRI tractography data acquired in healthy adults. When applying our trained model to the new data, neither FA nor streamline label is required. In this case, our method is robust to focal disruption or alteration of the FA due to the pathology. Fifth, considering that dMRI acquisition quality and the existence of brain lesions may affect the tracking of the RGVP fibers, we design an ensemble RGVP tractography method that enables highly sensitive tracking of RGVP for improved identification of the RGVP. This investigation extends our previous conference publication (Li et al., 2023) to include a retrospective analysis of an additional dataset from pituitary tumor patients, as well as an improved method description with in-depth technical details and an extensive experimental evaluation on data from additional sources.

The rest of this paper is organized as follows. Section 2 describes the datasets used, the proposed framework, and the model training and testing. Section 3 presents the experimental setup and results on high-quality dMRI data from healthy adults and typical clinical dMRI data from neurosurgical patients. Finally, the discussion, conclusions, and future work are given in Section 4.

## 2. METHODOLOGY

Our overall goal is to identify streamlines that belong to the RGVP from input tractography data generated in the skull base region. Figure 2 gives a visualization of example input tractography data, including RGVP streamlines selected using expert-drawn ROIs, and all other unselected non-RGVP streamlines resulting from false positive tracking or belonging to other structures. The RGVP and non-RGVP streamlines are visually highly similar in terms of their geometric trajectory; however, the non-RGVP streamlines fail to satisfy the strict anatomical ROI selection criteria. We can observe different FA values in local streamline regions of RGVP and non-RGVP streamlines (indicated using red arrows). This motivates the design of our microstructure-informed SCL (MicroSCL) method (Section 2.2.2). In addition, we can observe a highly imbalanced streamline distribution where the RGVP streamlines are fewer than the non-RGVP streamlines due to the use of a tractography seeding mask that is larger than the region that the RGVP passes through. This motivates the design of our streamline-level data augmentation (StreamDA) method to resolve any potential training biases due to the imbalanced input data (Section 2.2.4).

**Figure 2.**
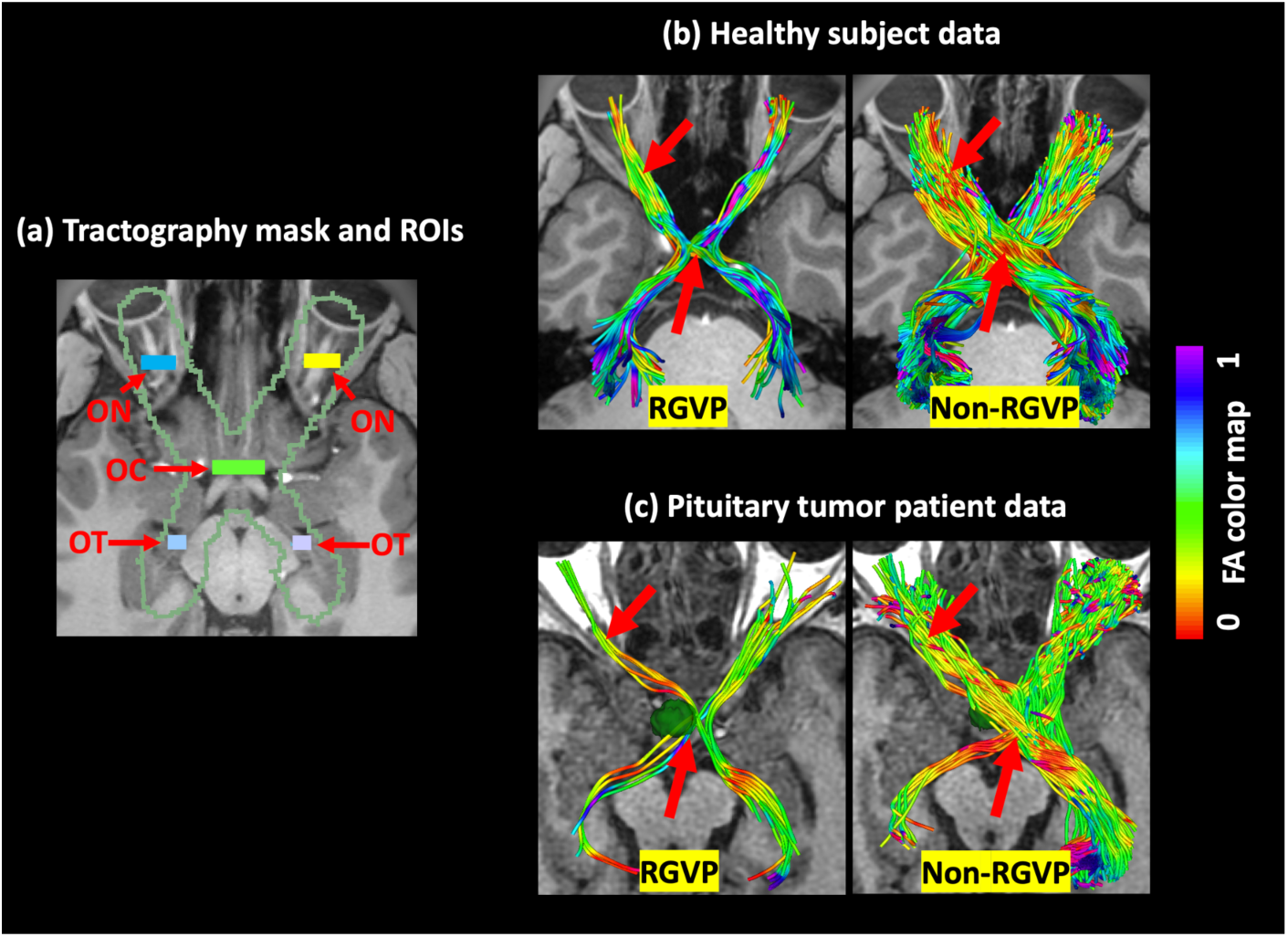
Example RGVP tractography data: (a) Tractography seeding mask and ROIs used for generating ground truth expert-selected data, (b) RGVP and non-RGVP streamlines of an example HCP subject, and (c) RGVP and non-RGVP streamlines of an example pituitary tumor patient. The red arrows in (b) and (c) show local streamline regions of RGVP and non-RGVP streamlines with different FA values.

### 2.1 Datasets

The study uses dMRI datasets from two different sources. The first dataset is from the Human Connectome Project (HCP) database (Van Essen et al., 2013) acquired from young healthy adults. The HCP dataset is used for model training and testing. The second dataset is acquired from patients with a diagnosis of pituitary tumor. The patient data is used for testing to demonstrate how DeepRGVP generalizes to data from a different source and a different population.

#### 2.1.1 HCP dataset

We use a total of 100 dMRI scans from the HCP database (Van Essen et al., 2013). The HCP data was acquired with a high-quality image acquisition protocol and preprocessed for data artifact correction (Glasser et al., 2013). The acquisition parameters were: TE/TR = 89.5/5520 ms, and voxel size = 1.25 × 1.25 × 1.25 mm^3^. In this study, we use the single-shell b = 1000 s/mm^2^ data because it has been shown to enable highly effective RGVP tracking (J. He et al., 2021). We perform a visual check of the dMRI data for each of the 100 subjects and exclude 38 subjects that had dMRI data with incomplete RGVP coverage due to face removal for data anonymization, as described in our previous work (J. He et al., 2021). Thus, tractography data from 62 subjects are used in this study.

Tractography is performed within a seeding mask (Figure 2(a)) using two-tensor unscented Kalman filter (UKF) tractography (Reddy & Rathi, 2016) via SlicerDMRI (Norton et al., 2017; Zhang, Noh, et al., 2020). We choose UKF because it can accurately track the RGVP (J. He et al., 2021) and other cranial nerves (Xie et al., 2020; Zhang, Xie, et al., 2020) and it allows the estimation of streamline-specific microstructure measures, including the measure of interest, FA. A streamline length threshold of 80 mm (a value lower than the length ∼100 mm of ground truth RGVP streamlines) is applied to eliminate any effect from streamlines too short to form the RGVP (Schiefer & Hart, 2007). In the UKF tractography method, the major parameters include *seedingFA*, *stoppingFA*, *Qm*, and *Ql*. Tractography is seeded in all voxels within the RGVP seeding mask, where FA is greater than the *seedingFA* threshold value. Tracking stops in voxels where the FA value falls below the *stoppingFA* threshold value. During the tracking, the UKF tractography methods use the *Qm* parameter to control process noise for angles/direction, and use the *Ql* parameter to control process noise for eigenvalues. Following our previous work that investigates how these major parameters affect the RGVP tracking (J. He et al., 2021), we set *SeedingFA* = 0.02, *stoppingFA* = 0.01, *Qm* = 0.001, *Ql* = 50, which was shown to generate the best RGVP reconstruction result.

We leverage ground truth RGVP streamlines selected using ROIs drawn by an expert (practicing neurosurgeon G.X.) and multi-rater validated in our previous study (J. He et al., 2021). These ROIs include the optic nerve (ON), optic chiasm (OC), and optic tract (OT) (Figure 2(a)). For model training, we also leverage the non-RGVP streamlines (Figure 2(b)) not selected by these ROIs. On average, there are ∼150 RGVP and ∼1200 non-RGVP streamlines per subject (thus, a ratio of ∼1:8 of RVGP to non-RGVP samples for training).

#### 2.1.2 Pituitary tumor patient dataset

We use dMRI from 10 pituitary tumor patients (PTPs) (see Table 1) from the First Affiliated Hospital of Sun Yat-sen University, China. All research procedures in this study were approved by the Ethics Committee of Clinical Research and Animal Trials of the First Affiliated Hospital of Sun Yat-sen University (Study Number [2022]057). The acquisition parameters are: TE = 97 ms, TR = 2700 ms, and voxel size = 1.5 × 1.5 × 1.8 mm^3^, with a field of view (FOV) of the skull base covering the RGVP. A total of 69 volumes were acquired for each subject, including 5 baseline images and 64 diffusion-weighted images at b = 1000 s/mm^3^.

**Table 1:** Demographics, diagnosis, and lesion size of the PTP dataset.

| *ID* | *Age (years)* | *Gender* | *Primary* | *Diagnosis* | *Lesion diameter* |
| --- | --- | --- | --- | --- | --- |
| 01 | 41 | Male | Yes | GH-PA | 8 mm |
| 02 | 56 | Female | Yes | NFPA | 17 mm |
| 03 | 44 | Male | Yes | GH-PA | 24 mm |
| 04 | 40 | Female | Yes | NFPA | 12 mm |
| 05 | 40 | Male | Yes | GH-PA | 9 mm |
| 06 | 34 | Female | Yes | Prolactinomas | 7 mm |
| 07 | 50 | Male | Recurrent | GH-PA | 4 mm |
| 08 | 37 | Male | Yes | GH-PA | 17 mm |
| 09 | 36 | Female | Yes | NFPA | 17 mm |
| 10 | 54 | Female | Recurrent | GH-PA | 15 mm |

Abbr: GH-PA: Growth hormone-secreting pituitary adenoma; NFPA: non-functioning pituitary adenoma; Prolactinomas: Prolactin-secreting adenoma.

Tractography of the RGVP in the PTP data is performed using the UKF algorithm, with a newly designed ensemble fiber tracking strategy. Unlike the high-quality HCP data, the PTP data was acquired with a protocol geared towards clinical applications with lower spatial and angular resolutions. RGVP fiber tracking is challenging in relatively low-quality data and is also impacted by the lesion affecting the RGVP. To increase the RGVP fiber tracking performance, we perform fiber tracking at various sets of tractography parameters and ensemble the tracked RGVP streamlines. This is inspired by the fact that tract-specific tracking parameters are needed for improved reconstruction of local fiber structures (Takemura et al., 2016; Tax et al., 2014; Wasserthal et al., 2019). In related work, ensemble fiber tracking has been used for effective RGVP reconstruction and evaluation of the decussation of optic nerves within the optic chiasm (Puzniak et al., 2019). In our study, we perform RGVP fiber tracking with varying combinations of *Qm* and *Ql*, as the tracking of specific RGVP segments is notably influenced by the settings of these parameters. Specifically, we compute 28 sets of RGVP tractography, with fiber tracking varying *Qm* across the range of 0.001 to 0.004 with a step size of 0.001 and *Ql* across the range of 50 to 350 with a step size of 50 (thus 4 *Qm* values and 7 *Ql* values). The *seedingFA* (=0.02) and *stoppingFA* (=0.01) are set to enable highly sensitive tracking of the RGVP (J. He et al., 2021). Then, all 28 sets of RGVP tractography results are assembled together as input to the proposed network.

To assess the RGVP identification results obtained with DeepRGVP, we perform RGVP streamline selection using expert-drawn ROIs, which are used as ground truth RGVP results. Unlike the HCP data where five ROIs (bilateral ONs, bilateral OTs, and the OC) are used, the OC ROI cannot be visually identified in the PTP dMRI data (see Supplementary Figure 1). Therefore, we draw only the ON and OT ROIs for the initial ROI selection, followed by the interactive selection of non-RGVP streamlines, where the expert (S.Y., a practicing neurosurgeon) interactively draws ROIs to exclude streamlines not belonging to the RGVP.

**Figure 3.**
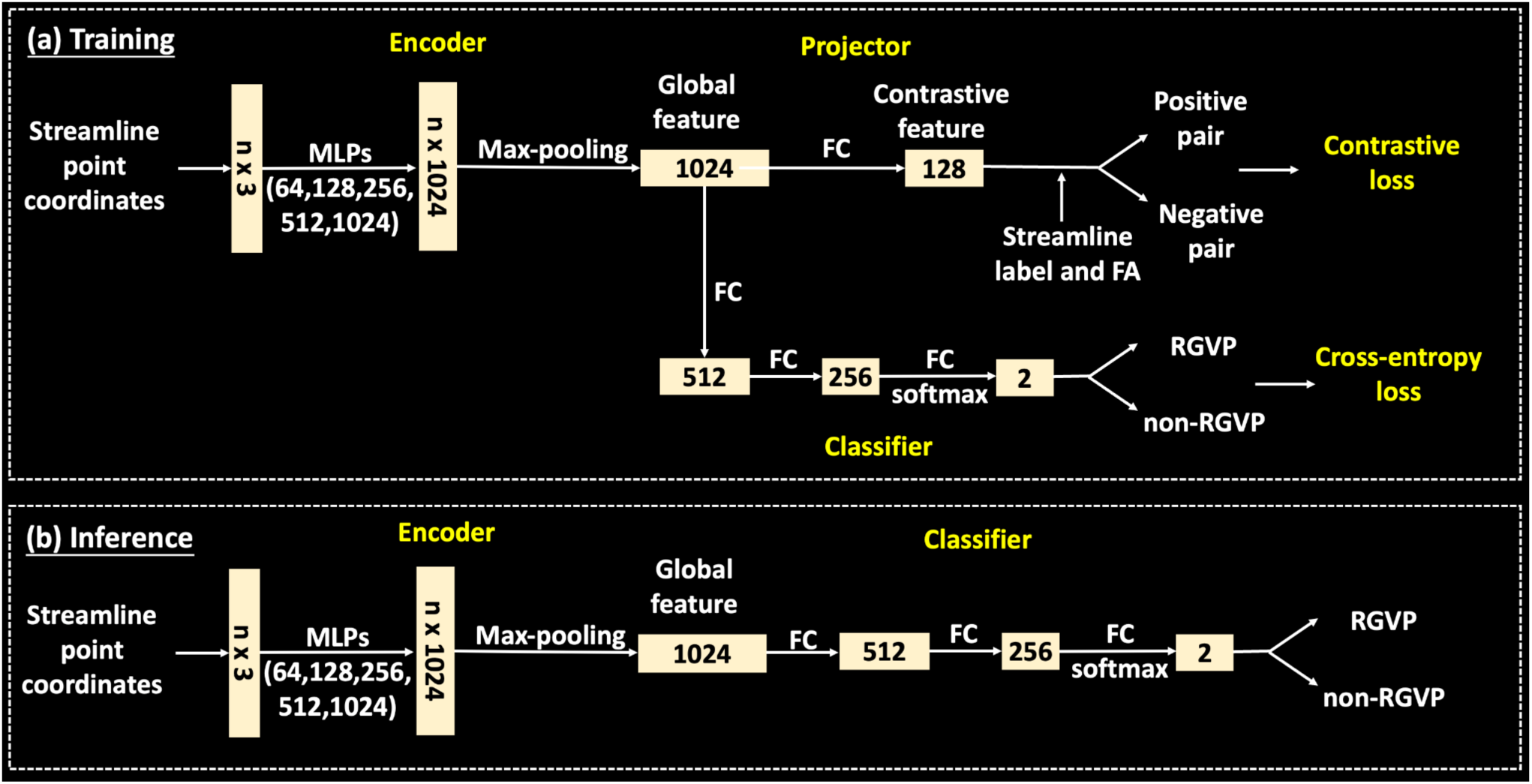
Overview of the DeepRGVP network, including the MicroSCL network and the downstream classifier for streamline classification.

### 2.2 Microstructure-informed SCL for automated RGVP identification

The overall framework of DeepRGVP includes the following major components. The first is a PointNet subnetwork that takes the feature representation of input tractography streamlines as point clouds and learns a global feature representation for each streamline (Section 2.2.1). The second component is a contrastive learning subnetwork that leverages both label information and tissue microstructure information to improve global features to be more discriminative between different classes (Section 2.2.2). The third component is a streamline classification subnetwork that classifies the input tractography streamlines into RGVP and non-RGVP based on the global features (Section 2.2.3). The fourth component is a streamline-level data augmentation process that increases the size of the training data to address the imbalanced training sample problem (Section 2.2.4).

#### 2.2.1 Streamline global feature learning via PointNet

The global feature representation is computed based on PointNet (Charles et al., 2017), which is widely used for point cloud classification and segmentation (Guo et al., 2020). Coordinates of tractography streamline points can be the input for PointNet networks, as used in (Astolfi et al., 2020; Y. Chen et al., 2023; Xue et al., 2023b) for tractography parcellation and filtering. An advantage of point cloud networks is that the global features generated are invariant to the order of points along a streamline, allowing the same global feature representation for forward and reverse point orders. The classic PointNet architecture includes a shared multi-layer perceptron (MLP), a symmetric aggregation function, and fully connected (FC) layers, as well as data-dependent transformation nets. In DeepRGVP, we make two modifications to the PointNet architecture by *1)* removing the transformation nets to preserve important information about the spatial position of streamlines (shown to be effective for tractography streamline parcellation in our previous work (Y. Chen et al., 2023; Xue et al., 2023a)) and *2)* adding additional MLP layers with sizes 256 and 512 between the layers of size 128 and 1024 (for a total of five layers of sizes 64, 128, 256, 512 and 1024) for better model training with the augmented data (see Section 2.2.4).

The detailed network architecture for learning the global feature is shown in Figure 3(a). The input of the encoder is RAS (Right, Anterior, Superior) point coordinates of a streamline; therefore, the dimension of the input is *n* × 3. We set *n* = 60, which is higher than *n* = 15 used in our previous work (Xue et al., 2023a) for the representation of the superficial white matter streamlines that generally have shorter lengths than the RGVP streamlines. Then, each point input is encoded by the 5 shared MLP layers, where each layer has batch normalization and rectified linear units (ReLU) activation, generating an output with *n* × 1024 dimensions. Following that, the global feature *g* of dimension 1 × 1024 is formed through max-pooling.

#### 2.2.2 Microstructure-informed supervised contrastive learning

Supervised contrastive learning (SCL) (Khosla et al., 2020) extends self-supervised contrastive learning (T. Chen et al., 2020) to a fully supervised mode by proposing a supervised contrastive loss, which aims to pull global features with the same class label closer in the latent space and push apart global features with different class labels. SCL has been shown to successfully improve the performance of supervised learning in computer vision (Khosla et al., 2020; Kopuklu et al., 2021; Schiffer et al., 2021; Zhong et al., 2021) and natural language processing (Gunel et al., 2020; Han et al., 2021; Huang et al., 2021) tasks, as well as in tractography parcellation tasks (Xue et al., 2023a). In DeepRGVP, we extend the SupWMA method (Xue et al., 2023a) by designing a microstructure-informed SCL (MicroSCL) method that leverages both streamline label (RGVP and non-RGVP) and tissue microstructure (FA) information to determine positive and negative pairs, rather than only relying on the label information as traditionally used in SCL. In this way, the learned global feature can better differentiate streamlines with similar trajectories but from different classes.

In detail, in addition to the traditional usage of label information for positive and negative sample pair determination, we include additional information about tissue microstructure to improve pair determination. To achieve this, we compute the absolute difference of mean streamline FA between each streamline pair. We constrain positive pairs to be streamlines from the same class that satisfy the following:

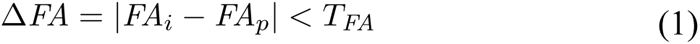

where *T_FA_* is a threshold on the allowable FA difference between streamlines in a positive pair.

Overall, the supervised contrastive loss *L_MicroSCL_* used in our study is:

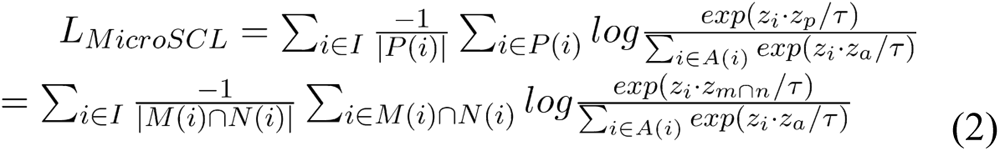

where *i* is a streamline belonging to a training batch *I*; *M*(*i*) is the set of streamlines with the same class label as streamline *i*; *N*(*i*) is the set of streamlines that satisfies the Δ*FA* condition in Eq (1); *P*(*i*) is the intersection of *M*(*i*) and *N*(*i*); *A*(*i*) is the set that includes all streamlines except for streamline *i* in batch *I*; *z_i_*, *z_p_* and *z_a_* are contrastive features of streamlines *i*, *p* ∈ *P*(*i*) and *a* ∈ *A*(*i*), respectively; τ (temperature) is a hyperparameter for optimization set to be 0.1 as suggested in (T. Chen et al., 2020).

#### 2.2.3 RGVP streamline classification

After obtaining the global feature for each streamline from MicroSCL, the streamline classification subnetwork predicts streamline class according to the global feature generated (Figure 3(a)). A simple subnetwork including 3 fully connected (FC) layers with sizes of 512, 256, and 2 (number of classes) and a cross-entropy loss is used.

#### 2.2.4 Streamline-level data augmentation

Data augmentation is a technique commonly used in deep learning to increase the diversity and quantity of training data by applying various transformations to the existing dataset (Shorten & Khoshgoftaar, 2019). This helps improve the generalization and robustness of machine learning models, particularly in scenarios where the original dataset is limited. Developing data augmentation methods to increase sample size is a known challenge in dMRI-related research (Barile et al., 2021). In our work, to curate a training dataset with a balanced sample distribution and an increased training sample size for improved model training, we propose the following StreamDA method. Each streamline consists of a sequence of points estimated by a tractography algorithm. For input to the network, we represent each streamline using *P* points sampled along the streamline. For data augmentation, we generate additional samples from each streamline by repeating the streamline point subsampling process multiple times, such that each time a different point subset with the same starting and ending points is generated (as demonstrated in Figure 4). We note that StreamDA is different from commonly used data augmentation strategies, such as data repetition and adding noise, where additional samples are synthetically generated. Since each point subset contains unique information, StreamDA can help the algorithm to learn the features of the RGVP and better differentiate RGVP from non-RGVP streamlines. In our study, given the original 1:8 ratio of RGVP to non-RGVP streamlines, we perform the StreamDA process 8 times for each RGVP streamline.

**Figure 4.**
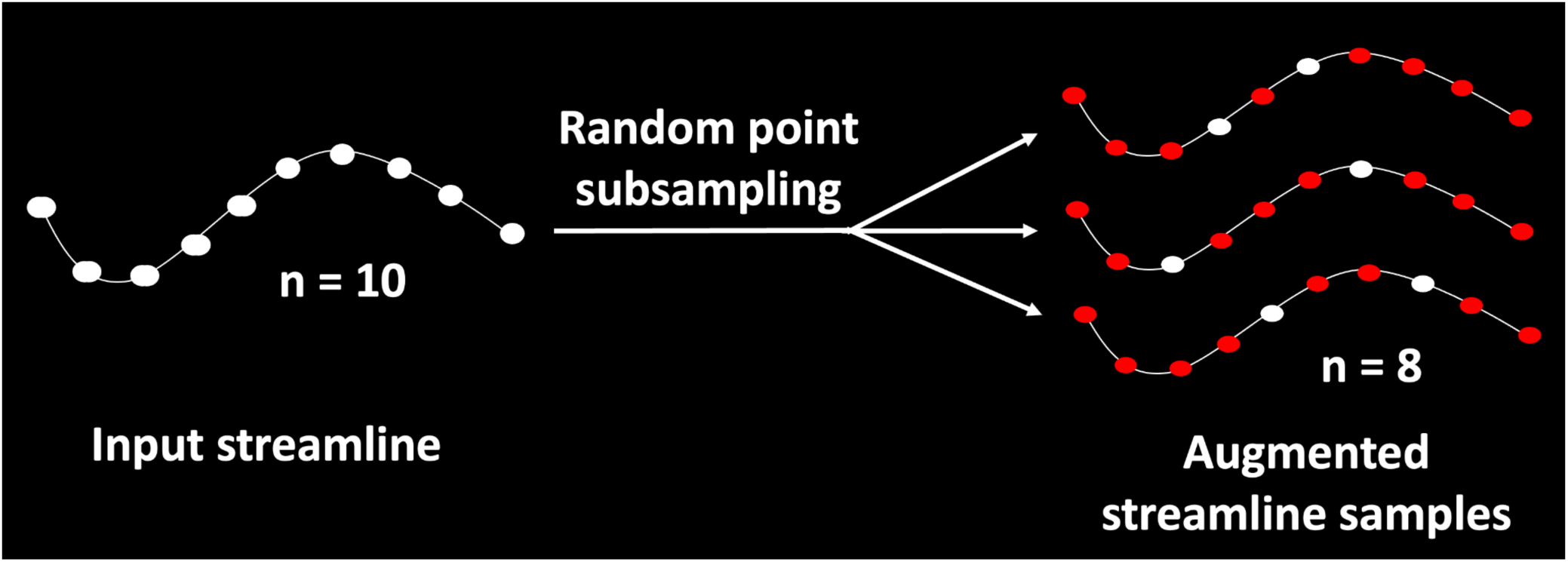
Graphic illustration of the proposed StreamDA method.

#### 2.2.5 RGVP Identification in new dMRI data

To identify the RGVP, the trained model is deployed to automate the identification process on new dMRI data with unlabeled tractography streamlines. The inference process works as follows (as illustrated in Figure 3(b)). First, the streamline point RAS coordinates of new dMRI tractography data are fed into the encoder subnetwork as the input, followed by a computation of the global feature for each streamline. Then, the generated global feature is fed into the classifier to predict the streamline labels (non-RGVP and RGVP streamlines) using the trained model. It is worth noting that during the training of the SCL model, both streamline labels and microstructure information are utilized. However, when applying the trained model to new data, neither of these is required.

#### 2.2.6 Implementation details

Our method is implemented using Pytorch (v1.7) (Paszke et al., 2019), and model training for HCP data is performed on an NVIDIA GeForce GTX 1080 Ti machine. All hyperparameters are set to the default settings suggested in SupWMA (Xue et al., 2023a), except for a modified learning rate (0.01) and batch size (512) that are tuned to accompany the addition of FA for pair determination. The threshold *T_FA_* in Eq (2) is set to be 0.1 (parameter search from 0.01 to 0.5). Both training phases utilize Adam (Kingma & Ba, 2014) as the optimizer with no weight decay. On average, each training epoch takes 4 seconds with 3GB GPU memory usage when using StreamDA.

### 2.3 Experimental evaluation

We perform the following experimental evaluation to assess the performance of DeepRGVP, including: 1) a comparison to three state-of-the-art methods for tractography parcellation (Section 2.3.1), 2) an ablation study to evaluate each DeepRGVP’s sub-component (Section 2.3.2), and 3) an assessment of DeepRGVP on pituitary patient data (Section 2.3.2).

#### 2.3.1 Comparison to state-of-the-art methods

We conduct a comparative analysis of DeepRGVP against three state-of-the-art (SOTA) deep learning approaches that classify streamlines into various categories for tractography parcellation within the white matter of the brain. These benchmark methods include *DeepWMA* (Zhang, Karayumak, et al., 2020), *DCNN* (H. Xu et al., 2019) and *SupWMA* (Xue et al., 2023a). In brief, DeepWMA (Zhang, Karayumak, et al., 2020) is designed for tractography streamline classification using a Convolutional Neural Network (CNN) and streamline spatial coordinate features. DeepWMA has eight CNN layers and three FC layers. It uses a “FiberMap” input feature that converts spatial coordinates of streamlines into 3-channel images that enable streamlines with forward and backward point orders to have nearly equivalent representations. DeepWMA achieves consistent deep white matter parcellation results across populations (Zhang, Karayumak, et al., 2020). DCNN (H. Xu et al., 2019) is also based on CNN for tractography streamline classification. DCNN was inspired by (K. He et al., 2015) and adapted in (H. Xu et al., 2018) to employ a soft spatial attention (ATT) module (K. Xu et al., 2015). During training, the DCNN method employs two losses: focal loss (Lin et al., 2017) to help in unbalanced datasets and center loss (CL) (Wen et al., 2016) to assist the network in obtaining better streamline representations. This framework obtains satisfactory accuracy for predicting functionally important white matter pathways that should be protected in the surgery of epilepsy patients (H. Xu et al., 2019). SupWMA (Xue et al., 2023a) is a point-cloud-based network (Qi et al., 2017) with supervised contrastive learning (SCL) (Khosla et al., 2020), which is designed for the classification of tract streamlines for superficial white matter parcellation. It has been shown to obtain highly consistent and accurate parcellation of superficial white matter across the lifespan in health and disease (Xue et al., 2023a). For each of the compared methods, we apply the suggested parameter settings described in the papers (Xue et al., 2023a; H. Xu et al., 2019; Zhang, Karayumak, et al., 2020) and the authors’ implementation. The comparison to the SOTA methods is performed using the HCP data. For each method, we train models using training data with and without the proposed StreamDA data augmentation. For each compared method and each HCP testing subject, the accuracy, F1, recall, and precision scores are computed between the prediction and the gold standard RGVP, and then the average score across all subjects is obtained.

#### 2.3.2 Ablation study

We perform an ablation study to evaluate the effects of the proposed MicroSCL and StreamDA. To do so, we compared the following methods. First, we compare with the *baseline* method that does not perform SCL (i.e., no supervised contrastive loss in our network) nor any data augmentation process. Second, we compare the *SCL_label_* method that performs label-based SCL, rather than the proposed microstructure-informed SCL. This method is performed on original training data without any data augmentation process. Third, we compare the *SCL_label+micro_* method that performs the proposed MicroSCL on original training data without any data augmentation process. Fourth, we compare with *SCL_label+micro_+Aug_repetition_* method that performs the proposed MicroSCL with DA by simple duplication of RGVP streamlines, rather than the proposed StreamDA. Fifth, we include the *SCL_label+micro_+Aug_StreamDA_*method that performs the proposed MicroSCL with StreamDA. The comparison to the SOTA methods is performed using the HCP data. For each compared method and each HCP testing subject, the accuracy, F1, recall, and precision scores are computed between the prediction and the gold standard RGVP, and then the average score across all subjects is obtained.

#### 2.3.3 RGVP identification in PTP data

We assess DeepRGVP’s generalization to dMRI data from neurosurgical patients with pituitary tumors. For each PTP dataset, we perform RGVP fiber tracking to obtain the input tractography data (as described in Section 2.1.2), followed by applying the trained DeepRGVP model. The model takes the input new tractography data and classifies them into non-RGVP streamlines and RGVP streamlines. For quantitative evaluation, in each patient dataset we compute the spatial overlap between the automated identification result and the expert-selected RGVP (Section 2.1.2). We employ the weighted Dice (wDice) score, an overlap metric used in many tractography parcellation studies (Cousineau et al., 2017; Zhang et al., 2019; Zhang, Karayumak, et al., 2020).

## 3. EXPERIMENTAL RESULTS

### 3.1 Comparison to state-of-the-art methods

As shown in Table 2, DeepRGVP generates the highest accuracy and the highest F1 using both original and augmented data, demonstrating the advantage of the proposed MicroSCL process. All methods except for DCNN achieve better performance using the augmented data compared to the original data, suggesting the benefit of our proposed DA strategy. In addition, DeepRGVP generates relatively high scores for both precision and recall, demonstrating balanced classification performance between RGVP and non-RGVP streamlines.

Figure 5 visually compares the identified RGVP across the compared SOTA methods. DCNN and DeepWMA are visually overinclusive with more streamlines compared to the expert-selected ground truth RGVP (e.g., see the inset images of Subjects 1 and 2). DeepRGVP and SupWMA are visually similar to expert selected RGVP. But DeepRGVP has improved sensitivity in identifying local structures, e.g., the ipsilateral pathway passing through the optic chiasm in Subject 2 (indicated by the red arrows).

**Table 2.**
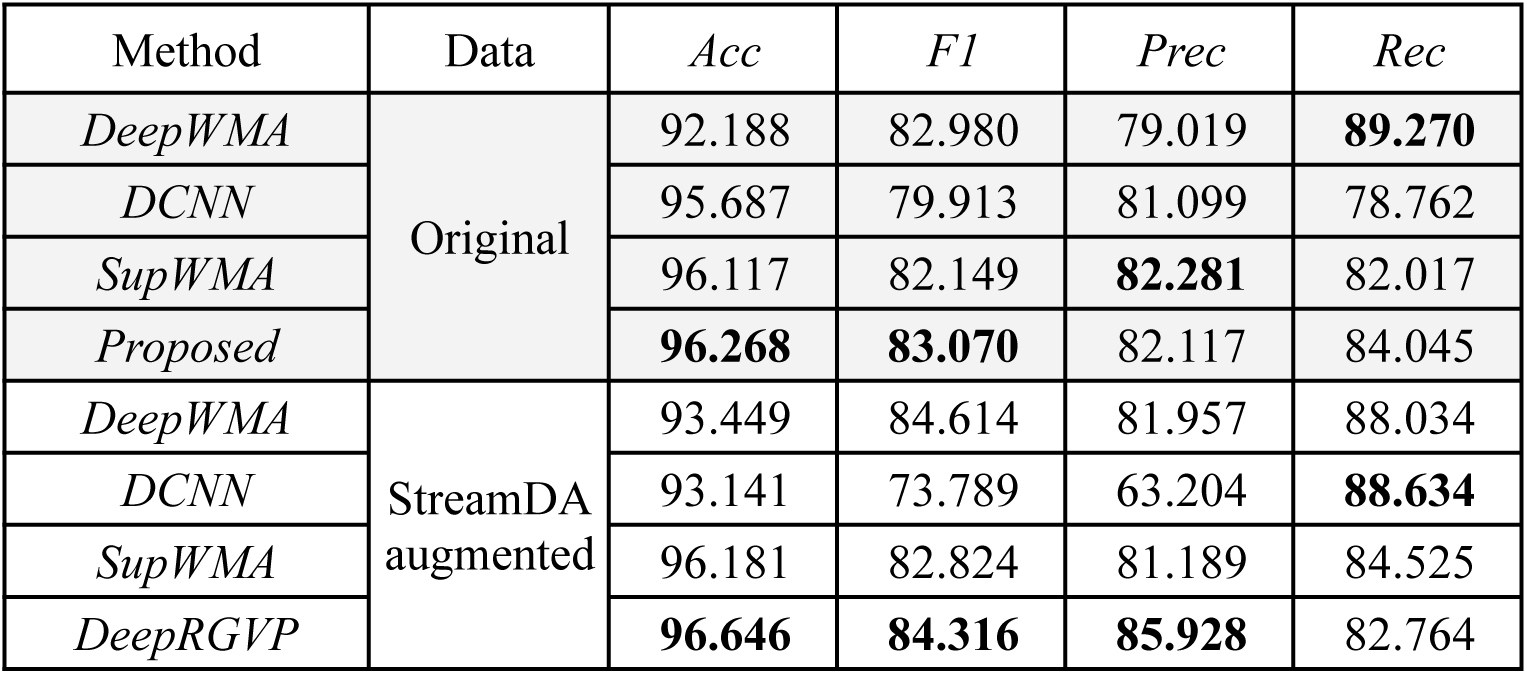
Quantitative comparisons with the state-of-the-art (SOTA) methods.

**Figure 5.**
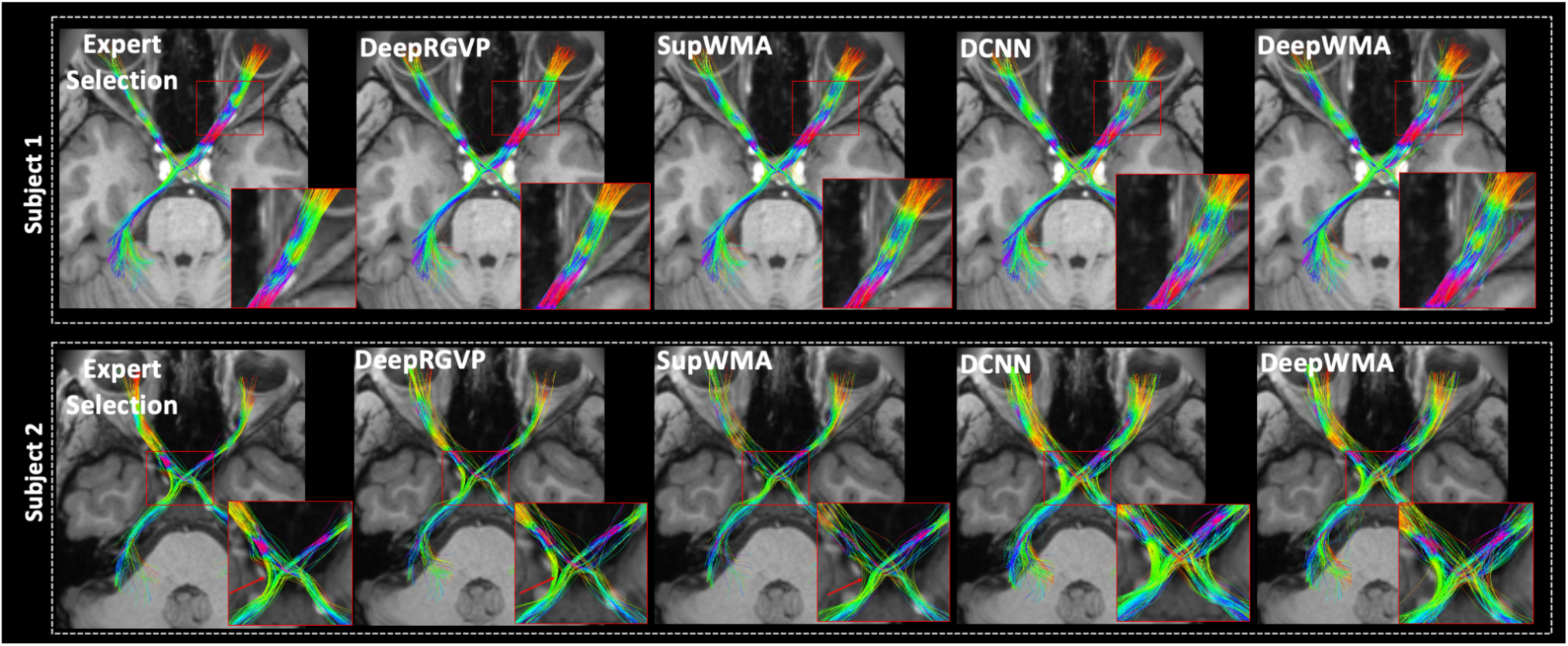
Visualization of expert selected and automatically identified RGVPs in two example HCP subjects. The inset zoomed images are provided for better visualization of local RGVP regions.

### 3.2 Ablation study

Table 3 provides the results of the ablation study. The higher accuracy and F1 scores obtained by *SCL_label_*compared with the baseline method (without SCL) show the benefit of using SCL for RGVP identification. The further improvement from *SCL_label+micro_*over *SCL_label_* shows the benefit of including microstructure information for better positive and negative sample pair selection in SCL. Furthermore, *SCL_label+micro_+Aug_repetition_*further improves the model performance compared to *SCL_label+micro_*, showing the advantage of using a balanced training dataset. However, the accuracy only slightly increases from 96.449% when using *SCL_label+micro_+Aug_repetition_* to 96.646% when using *SCL_label+micro_+Aug_StreamDA_*, while using the proposed *Aug_StreamDA_* can further improve the prediction accuracy and F1 scores. Overall, the proposed *SCL_label+micro_+Aug_StreamDA_*method achieves the best performance, showing the benefits of our network structure, in combination with the newly proposed microstructure-informed SCL and streamline-specific data augmentation, for improved RGVP identification.

**Table 3.** Results of the ablation study.

| Method | *Acc* | *F1* | *Prec* | *Rec* |
| --- | --- | --- | --- | --- |
| *Baseline* | 96.076 | 82.890 | 78.849 | **87.247** |
| $SCL_{label}$ | 96.117 | 82.149 | 82.281 | 82.017 |
| $SCL_{label+micro}$ | 96.268 | 83.070 | 82.117 | 84.045 |
| $SCL_{label+micro}+Aug_{repetition}$ | 96.449 | 83.410 | 84.909 | 81.964 |
| $SCL_{label+micro}+Aug_{StreamDA}$ | **96.646** | **84.316** | **85.928** | 82.764 |

### 3.3 RGVP identification in PTP data

Figure 6 shows a visualization of the identified RGVPs using the proposed DeepRGVP method from example PTP datasets. We can see that DeepRGVP enables successful RGVP identification in neurosurgical patients despite the effects of tumors affecting the RGVP fibers. In addition, the identified RGVP is highly visually comparable to the expert selected RGVPs. Our quantitative evaluation also shows a high spatial overlap between RGVPs obtained from the proposed method and the expert selection, with a high wDice = 0.82 ± 0.08 (a wDice score over 0.72 is suggested to be a good tract overlap (Cousineau et al., 2017; Zhang et al., 2019)).

**Figure 6.**
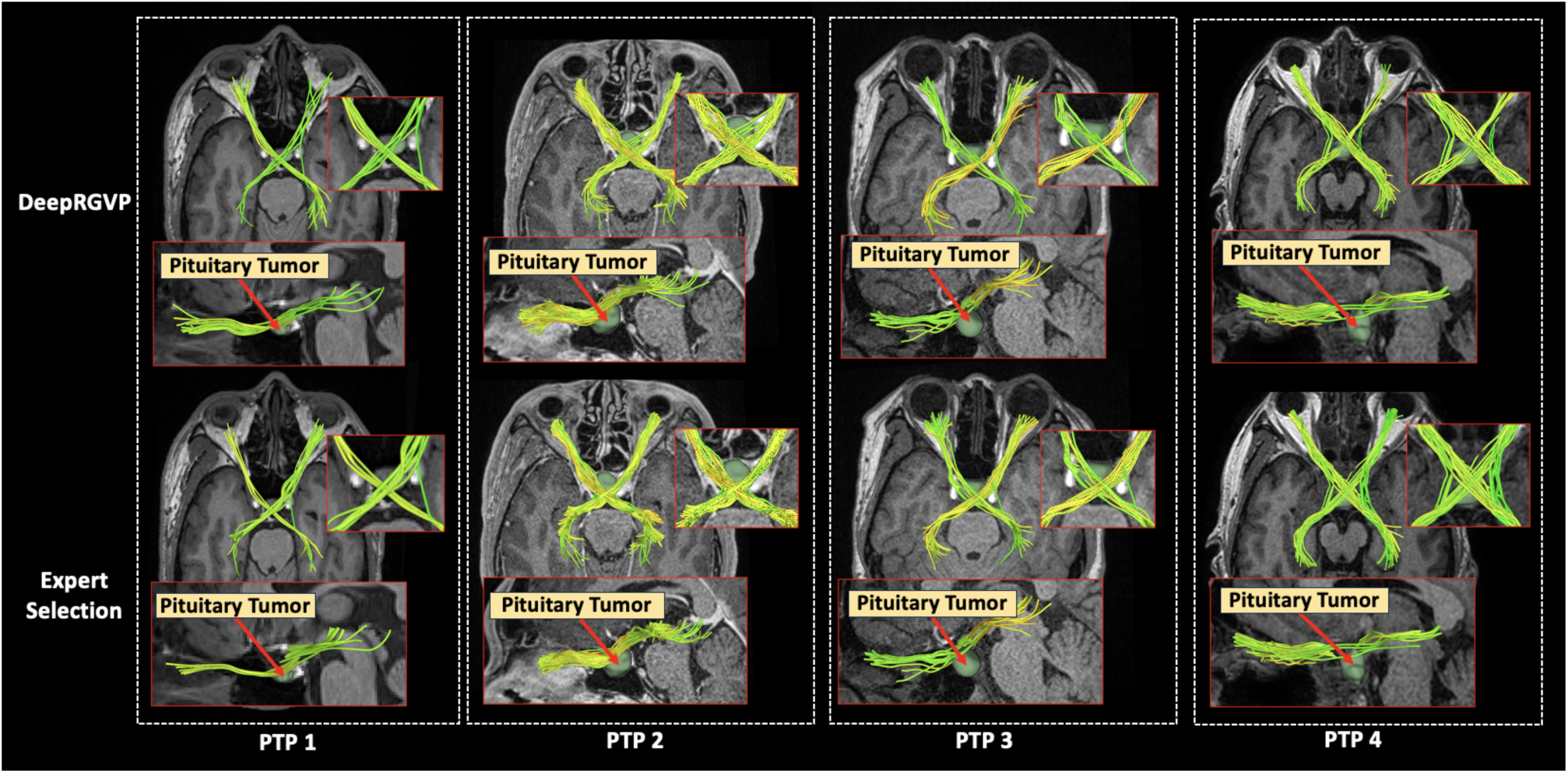
Visual comparison of the identified RGVPs in four example neurosurgical patients with ground truth expert-selected RGVP. The inset images are provided for better visualization of local RGVP regions.

## 4. DISCUSSION

In this work, we present a novel deep-learning framework to enable automated identification of the RGVP. We design a novel microstructure-informed supervised contrastive learning method that leverages both streamline label and tissue microstructure information to determine positive and negative pairs. We propose a simple and successful streamline-level data augmentation method to address highly imbalanced training data. We perform comparisons with several state-of-the-art deep learning methods designed for tractography parcellation, and we show superior RGVP identification results using DeepRGVP. We also demonstrate a successful application of DeepRGVP in neurosurgical patient data with pituitary lesions affecting the RGVP.

We demonstrate that DeepRGVP outperforms several state-of-the-art methods. The DeepWMA and DCNN methods yield less favorable RGVP identification results than DeepRGVP and SupWMA, in terms of quantitative metrics and visualization. This is likely due to the fact that DeepWMA and DCNN are based on traditional CNNs with input in the form of feature vectors or 2D images, while DeepRGVP and SupWMA use an advanced SCL framework with point clouds for streamline representation. Compared to SupWMA, the proposed SupWMA achieves improved identification accuracy and F1 scores. This is likely due to the novel design of the microstructure-informed SCL strategy that can learn a more discriminative global feature for the classification of RGVP and non-RGVP streamlines.

The proposed streamline-level data augmentation, StreamDA, can help improve the identification of the RGVP. Across the tractography parcellation methods under study, the DeepWMA, SupWMA, and DeepRGVP methods all benefited from the data augmentation, leading to improved RGVP identification accuracy compared to using the original unbalanced training data. However, we noticed a decreased performance of DCNN when using the augmented (balanced) data. This may relate to the design of the DCNN method to perform well on unbalanced datasets (H. Xu et al., 2019). This suggests that data augmentation should be used with careful consideration of the design of the classifier.

We demonstrate the generalization of DeepRGVP to dMRI data acquired using different imaging protocols. The ability of a dMRI parcellation method to generalize to data from different acquisitions is important, because there is no standard, and dMRI acquisitions can have widely varying properties such as spatial and angular resolutions, posing a challenge for machine learning approaches. In our study, we train a deep learning model from the high-quality HCP data, from which the reconstruction of RGVP fibers is relatively easy due to the high spatial and angular resolutions. For the data acquired with a clinical acquisition protocol, we design an ensemble RGVP fiber tracking strategy to improve RGVP fiber tracking so that the RGVP fibers can be effectively reconstructed. In this case, the DeepRGVP method can successfully identify the RGVP by classifying the RGVP and non-RGVP streamlines.

We show that our method enables automated identification of the RGVPs in pituitary tumor patients. Despite any potential effects from lesions affecting the RGVP, we show good RGVP identification generalization performance in the patient data, even though the RGVP identification model is trained using data from healthy subjects. In related work, previous studies have shown successful parcellation of the major long-range white matter tracts such as the arcuate fasciculus and corticospinal tract in patient data (Wasserthal, Jakob, Peter Neher, and Klaus H. Maier-Hein., 2018; Zhang et al., 2018; Zhang, Karayumak, et al., 2020). To the best of our knowledge, our method is the first to show successful automated identification of the RGVP in neurosurgical patient data. This can improve the current manual selection strategy to resolve inter-observer variability and may improve clinical efficiency, possibly serving as an important tool for surgical planning in diseases affecting the RGVP, e.g. pituitary tumors.

Potential limitations of the present study, including suggested future work to address limitations, are as follows. First, while our method is designed for the identification of the RGVP, it can generally be used to identify any other cranial nerves and/or anatomical fiber tracts from tractography data. It would be an interesting future investigation to test the performance of DeepRGVP on these brain structures. Second, we demonstrated the generalization of DeepRGVP on pituitary patient data, where the RGVP is mostly affected around the optic chiasm region. A future investigation could include an evaluation of RGVP identification in the setting of other diseases affecting the RGVP (such as optic neuritis, optic gliomas, optic nerve sheath meningiomas, and others), where different regions of RGVP might be affected due to inflammation and tumor infiltration, rather than displacement and tract compression. Third, when generalizing to new patient data, we showed the proposed method obtained highly comparable results to the expert manual selection results, but we noticed that both methods may fail to identify certain subdivisions of the RGVP, e.g., the non-decussating RGVP portion. This was related to the fact that the relatively low imaging quality and effects of lesions influenced the RGVP tracking performance. Future work could include designing an RGVP-specific tractography method to enable improved RGVP reconstruction. Fourth, one of the benefits of our method is that the identification of RGVP in new tractography data only uses the streamline trajectory information but not the microstructure FA information. It would be interesting to investigate whether including such information can help with global feature computation in new data. However, this would require harmonization of the testing dMRI data with the training data to remove scanner-specific differences, considering the substantial variability in microstructure parameters computed from dMRI data from various sources (Cetin Karayumak et al., 2019). Also, this would require considering the situation where there is a focal disruption or alteration of tissue microstructure property due to the pathology. In contrast, the existing approach remains generally applicable to diverse tractography data from multiple acquisitions.

## 5. CONCLUSION

In this paper, we present a novel deep learning method, *DeepRGVP*, to enable automated identification of the RGVP using diffusion MRI tractography. Experimental results show that DeepRGVP achieves improved identification accuracy compared to several state-of-the-art methods. In addition, we show a successful application of DeepRGVP to neurosurgical patient dMRI data acquired with different MRI acquisition protocols. We anticipate that DeepRGVP will be a useful tool in dMRI-based analyses of the RGVP, e.g., in neurosurgical planning research. Overall, our study shows the high potential of using deep learning to automatically identify the RGVP.

## 6. DATA AND CODE AVAILABILITY

The data used in this project include the public Human Connectome Project (HCP) dataset (www.humanconnectome.org). The PTP data is our in-house dataset, which is not publicly available because public availability would compromise participant confidentiality and participant privacy. The code and trained model will be available at: https://github.com/SlicerDMRI/DeepRGVP.

## ACKNOWLEDGEMENTS

This work is in part supported by the National Key R&D Program of China (No. 2023YFE0118600), the National Natural Science Foundation of China (No. 62371107, No. 62303413), the Guangdong Basic and Applied Basic Research Foundation (2023A1515011644), and the National Institutes of Health (R01MH108574, P41EB015902, R01MH074794, R01MH125860, R01MH119222, R01MH132610, R01NS125781). F.Z. acknowledges a BWH Radiology Research Pilot Grant Award.

---

**Supplementary Figure 1:**
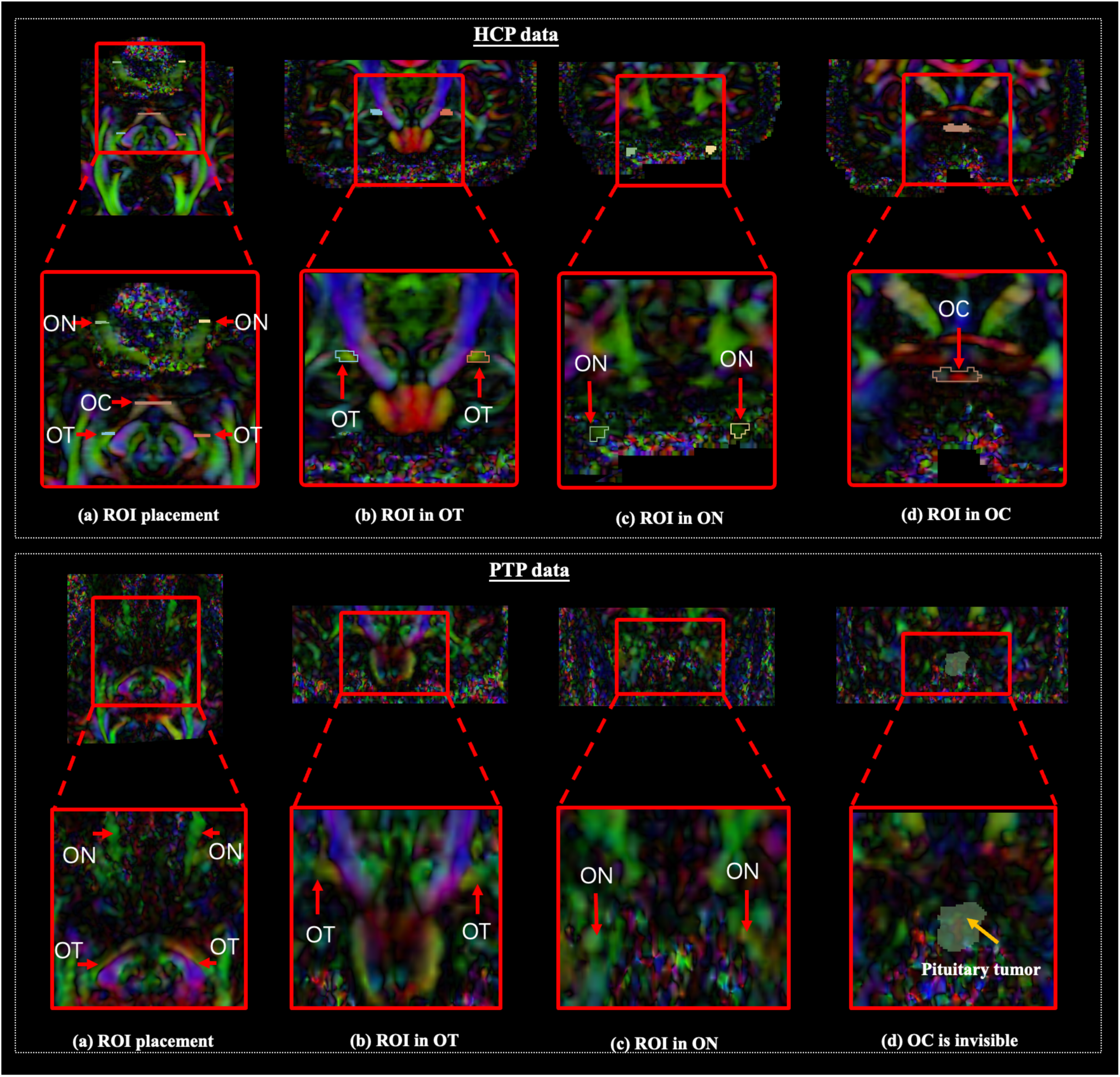
The top subfigure shows the placement of the ON, OT and OC ROIs in one example HCP subject. The bottom subfigure shows the ROI placement of the OTs and ONs, while the OC ROI can not be visually identified possibly due to effects of the pituitary tumor.

## Notes

### Competing Interest Statement

The authors have declared no competing interest.

